# CoV-AbDab: the Coronavirus Antibody Database

**DOI:** 10.1101/2020.05.15.077313

**Authors:** Matthew I. J. Raybould, Aleksandr Kovaltsuk, Claire Marks, Charlotte M. Deane

## Abstract

The emergence of a novel strain of betacoronavirus, SARS-CoV-2, has led to a pandemic that has been associated with hundreds of thousands of deaths. Research is ongoing around the world to create vaccines and therapies to minimise rates of disease spread and mortality. Crucial to these efforts are molecular characterisations of neutralising antibodies to SARS-CoV-2. Such antibodies would be valuable for measuring vaccine efficacy, diagnosing exposure, and developing effective biotherapeutics. Here, we describe our new database, CoV-AbDab, which already contains data on over 380 published/patented antibodies and nanobodies known to bind to at least one betacoronavirus. This database is the first consolidation of antibodies known to bind SARS-CoV-2 and other betacoronaviruses such as SARS-CoV-1 and MERS-CoV. We supply relevant metadata such as evidence of cross-neutralisation, antibody/nanobody origin, full variable domain sequence (where available) and germline assignments, epitope region, links to relevant PDB entries, homology models, and source literature. Our preliminary analysis exemplifies a spectrum of potential applications for the database, including identifying characteristic germline usage biases in receptor-binding domain antibodies and contextualising the diagnostic value of the SARS-CoV binding CDRH3s through comparison to over 500 million antibody sequences from SARS-CoV serologically naive individuals. Community submissions are invited to ensure CoV-AbDab is efficiently updated with the growing body of data analysing SARS-CoV-2. CoV-AbDab is freely available and downloadable on our website at *http://opig.stats.ox.ac.uk/webapps/coronavirus*.

## Introduction

To respond effectively to the recent Severe Acute Respiratory Syndrome Coronavirus 2 (SARS-CoV-2) pandemic, it is essential to understand the molecular basis for a successful immune response to coronavirus infection (1). In particular, characterising the B-cell response is important as the identification of potent neutralising antibodies could pave the way for effective treatments, aid in prior exposure diagnosis, or assist in predicting vaccine efficacy (2–5).

Molecular characterisations of binding/neutralising antibodies to SARS-CoV-2 antigens are only just beginning to emerge. However, the SARS-CoV-2 and SARS-CoV-1 (the virus responsible for the 2003 epidemic) spike protein receptor binding domains (RBDs) target the same human receptor and share high sequence and structural homology (2). As a result, collating data on SARS-CoV-1 binders may lead to the identification of potent cross-neutralising antibodies, as suggested in some early SARS-CoV-2 studies (6, 7). Solved crystal and cryo-EM structures indicate a relatively discrete set of neutralising RBD epitopes (possibly resulting from substantial glycan coverage (8)), with paratopes tending to span both the heavy and light chain complementarity-determining regions (6, 9–12).

Other SARS-CoV-2 surface proteins also display homology to more distantly related betacoronaviruses such as the Middle East Respiratory Syndrome coronavirus (MERS-CoV). Therefore, knowledge of antibodies that bind to MERS-CoV antigens could be relevant in treating SARS-CoV-2 infection, and indeed the anti-MERS-CoV combination therapy REGN3048/REGN3051 is already being trialled on SARS-CoV-2 patients in the USA (13).

Given this, a central database facilitating molecular-level comparisons between published and patented anti-coronavirus antibodies would be a valuable tool in the fight against COVID19. This resource would also act as a central hub to consolidate knowledge and coordinate efforts to identify novel antibodies that neutralise SARS-CoV-2. As the number of known binders builds up over time, researchers could harness this repository for many purposes, including deriving crucial sequence/structural patterns that distinguish neutralising from non-neutralising SARS-CoV-2 binders (1), or deducing independent neutralising epitopes exploitable by combination therapies.

We have built CoV-AbDab, a new database that aims to document molecular information and metadata on all published or patented anti-coronavirus antibodies.

## Data Sources

Academic papers and patents containing coronavirus-binding antibodies were primarily sourced by querying PubMed, BioRxiv, MedRxiv, GenBank, and Google Patents with relevant search terms. Several review articles were helpful in ensuring maximal coverage, in particular those by Coughlin and Prabhakar (14), Du *et al*. (15), Zhou *et al*. (16), Shanmugaraj *et al*. (17), and Jiang *et al*. (18). If the variable domain sequence was available, ANARCI (19) was used to number sequences in the IMGT (20) numbering scheme, and to assign V and J gene origins. In some cases we could source germline assignments and/or CDR3 sequences from the source literature for antibodies where the full Fv sequence was not supplied. Our Structural Antibody Database (21), which tracks all antibody structures submitted to the Protein Data Bank (22) (PDB), was mined to identify relevant solved structures. Our antibody/nanobody homology modelling tool, ABody-Builder (23), was used to generate full Fv region structural models where no solved structures were available.

## Contents

CoV-AbDab is an effort to document all coronavirus binding/neutralising antibodies and nanobodies reported in academic publications and commercial patents. Where possible, the following information is documented for each entry:

1. The published name of the antibody/nanobody
2. Antigens that the antibody/nanobody has been proven to bind and/or neutralise.
3. The protein domain targeted by the antibody/nanobody (e.g. spike protein receptor binding domain)
4. The developmental origin of the antibody/nanobody (e.g. engineered/naturally raised, species information, *etc*.)
5. Sequence information including: (a) the entire variable domain sequence for the antibody/nanobody, highlighting the CDR3 regions, and (b) V and J gene germline assignments.
6. Links to any available structures involving the antibody/nanobody
7. (If Fv sequence available) A homology model of the antibody/nanobody
8. References to the primary literature on the antibody/nanobody
9. Timestamps to show when the antibody/nanobody was added and last updated
10. Any steps we are taking to follow up on the entry (e.g. to source its sequence and/or add further metadata)

As of 14^th^ May 2020, CoV-AbDab contains 385 entries across 46 publications (6, 7, 9, 11, 12, 24–64) and 19 patents. Of these, 156 entries are associated with MERS-CoV, 149 are associated with SARS-CoV-1, and 105 are associated with SARS-CoV-2 (each entry may be tested against multiple coronaviruses). It lists 263 unique full variable domain antibody/nanobody sequences and 56 links to relevant PDB structures, which include coronavirus spike proteins bound to their native receptors (35, 65–72). We are continuing to contact authors to confirm whether missing sequences can be recovered and added to existing entries. If sequences have been lost or cannot be released, they have been removed from the database and confirmed as such in a separate list on the CoV-AbDab homepage.

## Analysis

The following analysis was carried out on the CoV-AbDab database as of 10^th^ May 2020. For clarity, we use the term “SARS-CoV-1” to refer specifically to the virus that caused the 2003 epidemic, and “SARS-CoV” to refer to binders to SARS coronaviruses in a general sense.

### Developmental Origins and Targets

We first analysed the developmental origins of antibody/nanobody binders to SARS CoV-1/2 (Figure 1a) and MERS-CoV (Supplementary Figure 1a). The vast majority of the SARS-CoV antibody binders have human genetic origin (88.5% with sequence information aligned to human germlines), and derive from a mixture of isolated B-cells from infected or convalescent patients, transgenic mice, or recombinant human immune or non-immune phage display libraries. We soon expect the proportion from infected human B cells to increase, as papers characterising and panning the adaptive immune responses of SARS-CoV-2 patients continue to emerge (24–26). A relatively small portion of antibodies were detected by challenging mice with SARS antigens, and a few of these were subsequently humanised. All but one SARS-CoV binding nanobody was obtained using phage display. MERS-CoV antibodies followed a similar distribution of origins, but nanobodies were sourced from the B-cells of infected/convalescent camels or immunised llamas (Supplementary Figure 1b).

**Fig. 1.**
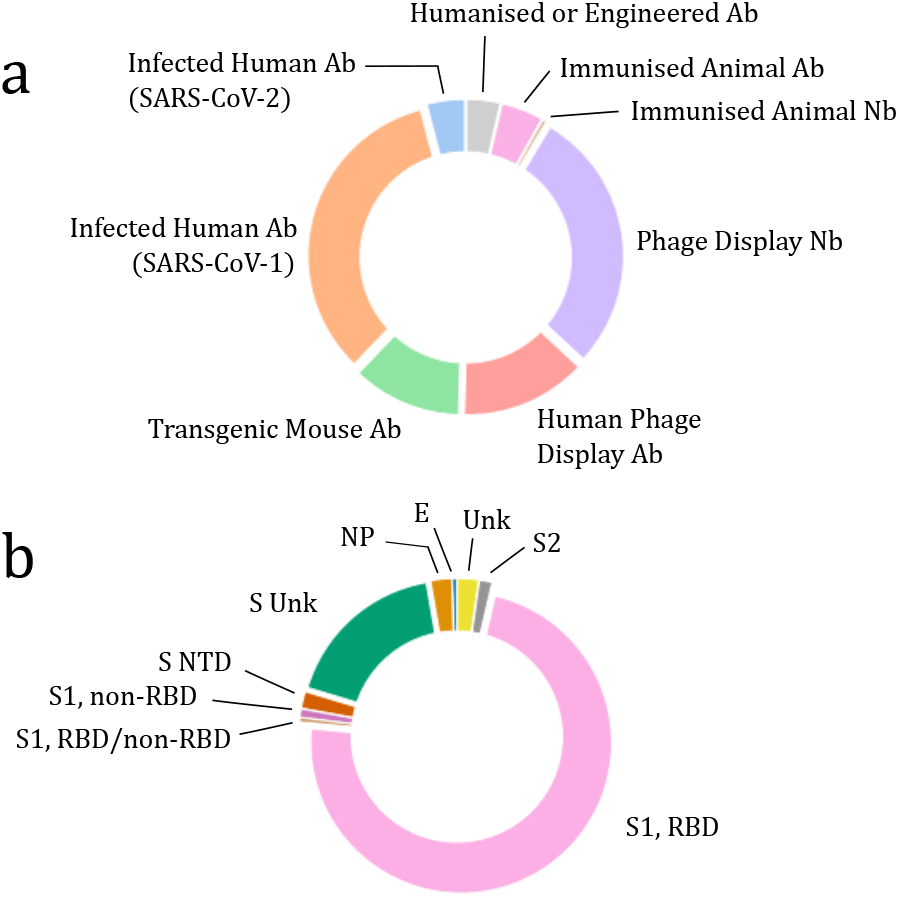
Donut charts showing (a) the origins of all identified SARS-CoV-1/2 binders and (b) the protein targets of identified SARS-CoV-1/2 binders. Spike protein binders are further classified by targeted domain. Equivalent plots for MERS-CoV binders are available in Supplementary Figure 1. S = Spike protein, NP = Nucleocapsid protein, E = Envelope protein, Unk = Unknown, NTD = N-Terminal Domain, RBD = Receptor Binding Domain, S1 = Spike protein S1 domain, S2 = Spike protein S2 domain.

We also evaluated the distribution of protein targets (and epitope regions, for spike protein binders) for all anti-SARS-CoV1/2 (Figure 1b) and anti-MERS-CoV (Supplementary Figure 1b) antibodies/nanobodies. The spike (S) protein is known to mediate coronaviral entry into cells through a biochemical signal initiated by RBD-ACE2 (SARS-CoV) or RBD-DPP4 (MERS-CoV) binding (65, 71). Therefore, antibodies/nanobodies that can attach to this domain are of particular pharmacological interest, as they may block a crucial step of the viral reproductive cycle, neutralising the infection. This bias was strongly reflected in the observed coronavirus protein targets, with 72.9% of SARS-CoV binders and 58.3% of MERS-CoV binders attacking the spike protein RBD. A few other S protein domains were represented, such as the S2 domain and N-Terminal Domain, as well as some binders to the nucleocapsid and envelope proteins.

At the time of writing, sequence information has been released for three antibodies (CR3022, S309, and S315) and one nanobody (VHH-72) that have been proven to neutralise SARS-CoV-2. These all target the RBD, and can cross-neutralise SARS-CoV-1.

### Genetic Origins

In constructing our database, we evaluated/collected the gene transcript origins of as many of the anti-SARS-CoV and anti-MERS-CoV antibodies as possible. Here, we analyse IGHV gene usage, as this transcript encodes two of the three heavy chain complementarity determining regions (CDRH1 and CDRH2). Analysis of the CDRH3 region, which lies at the junction of IGHV, IGHD, and IGHJ genes, is performed in the next section.

Figure 2a shows the distribution of IGHV genes in SARS-CoV binding antibodies against all targets (left-hand-side), and after filtering only for antibodies known to bind the RBD (right-hand-side). In both cases, over half of the antibodies compromise one of four V genes: IGHV3-30, IGHV1-2, IGHV3-23, and IGHV1-18. The dominant V gene, IGHV3-30, was identified in spike protein binders from five independent investigations — Pinto *et al*. (6), Sui *et al*. (34), Hwang *et al*. (35), and patents WO2008060331A2 and CN1903878A — and is present in around 21% of RBD binders (monoclonal antibody 80R and its variants are counted as a single source). IGHV3-30 has been found to be unusually abundant in several recent B-cell sequencing investigations (26, 74–76). IGHV1-2 (75, 76) and IGHV1-18 (76) have also been implicated.

**Fig. 2.**
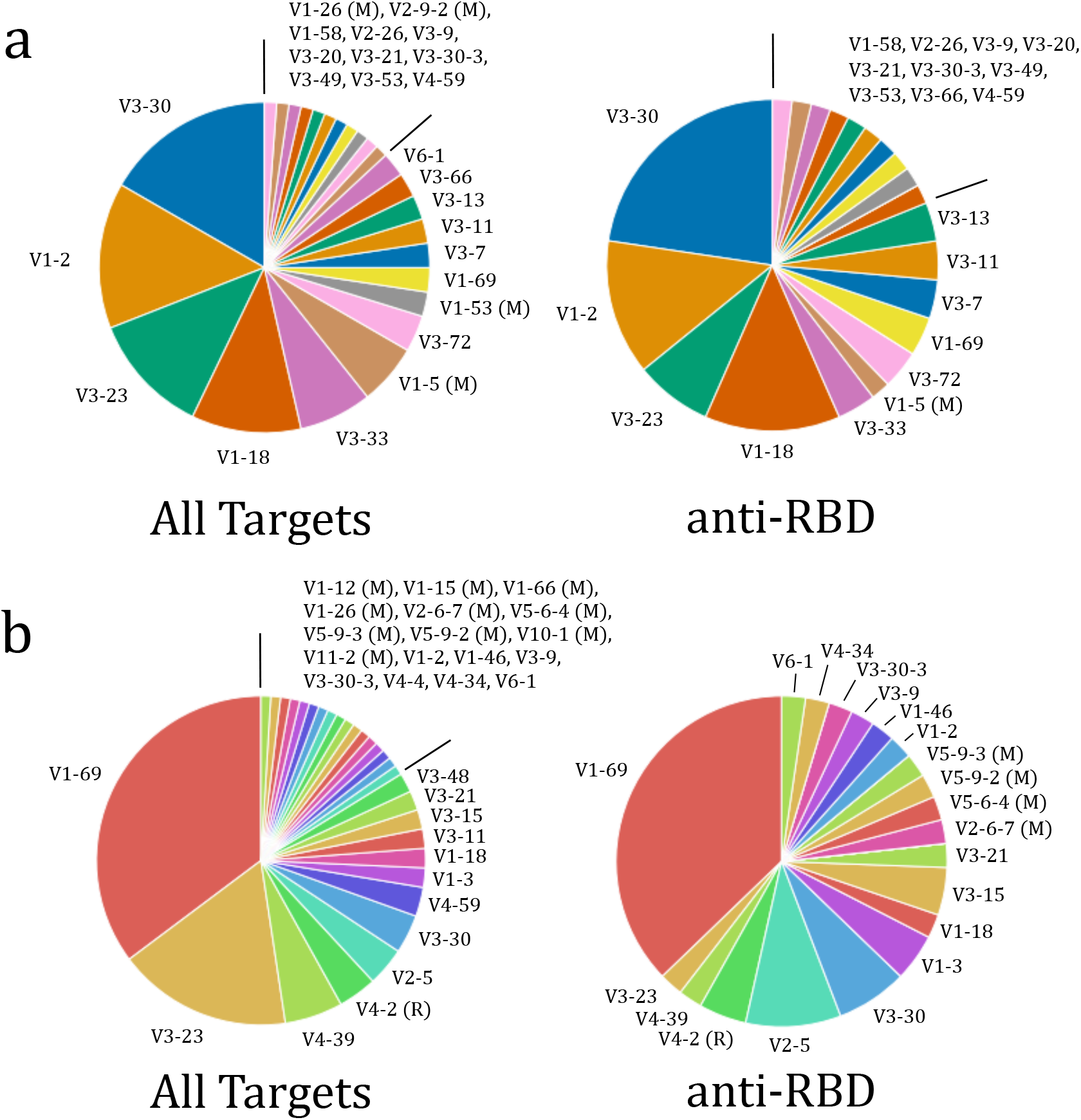
Pie charts showing the distributions of IGHV gene usage in (a, LHS) SARS-CoV binding antibodies, (a, RHS) SARS-CoV Receptor Binding Domain (RBD) binding antibodies, (b, LHS) MERS-CoV binding antibodies, and (b, RHS) MERS-CoV RBD binding antibodies. Monoclonal antibody 80R and its five closely-related variants were counted as a single entry in both SARS plots. All germlines are human, unless appended with M (Murine) or R (Rhesus).

In marked contrast to the SARS data, anti-MERS-CoV RBD antibodies (Figure 2b) are disproportionately (37.2%) sourced from the IGHV1-69 locus. These antibodies derive from eight independent investigations — Wang *et al*. (47), Niu *et al*. (48), Chen *et al*. (51), Ying *et al*. (60), Jiang *et al*. (61), Tang *et al*. (62), and patents WO2015179535 and WO2019039891. The IGHV1-69 transcript is commonly observed in broadly neutralising antibody responses, for example to the influenza hemagglutinin stem domain (77).

### CDRH3 Analysis

We then analysed the CDRH3 regions of anti-SARS-CoV and anti-MERS-CoV antibodies. Overall, we traced 54 distinct SARS-CoV RBD binding antibody CDRH3 sequences and 75 distinct MERS-CoV RBD binding antibody CDRH3 sequences. The non-redundant CDRH3 length distributions are shown in Supplementary Figure 2. SARS-CoV RBD binders are spread between CDRH3 lengths 8 and 20 (median: 16, mean: 14.87 ± 3.56), while the CDRH3s of MERS-CoV RBD binders lie between lengths 5 and 26 (median: 18, mean: 16.17 ± 4.14). The longer average lengths for MERS-CoV binders are consistent with the observed IGHV gene distribution, as broadly neutralising IGHV1-69 antibodies tend to have longer CDRH3s (21).

To see whether RBD binding CDRH3s displayed any sequence biases, we used WebLogo plots (73) to visualise residue/position distributions (Figure 3). The MERS-CoV RBD binders displayed slightly higher homology at central loop positions, but neither showed a strong signal that implicates a particular interaction type. The SARS-CoV RBD binders have a slight tendency to exploit a poly-tyrosine tail towards the end of the CDRH3, hinting at a role for the IGHJ6 germline that bears this motif. IGHJ6 was independently implicated in a clone convergent in four of six SARS-CoV-2 patients in the study by Nielsen *et al*. (26).

**Fig. 3.**
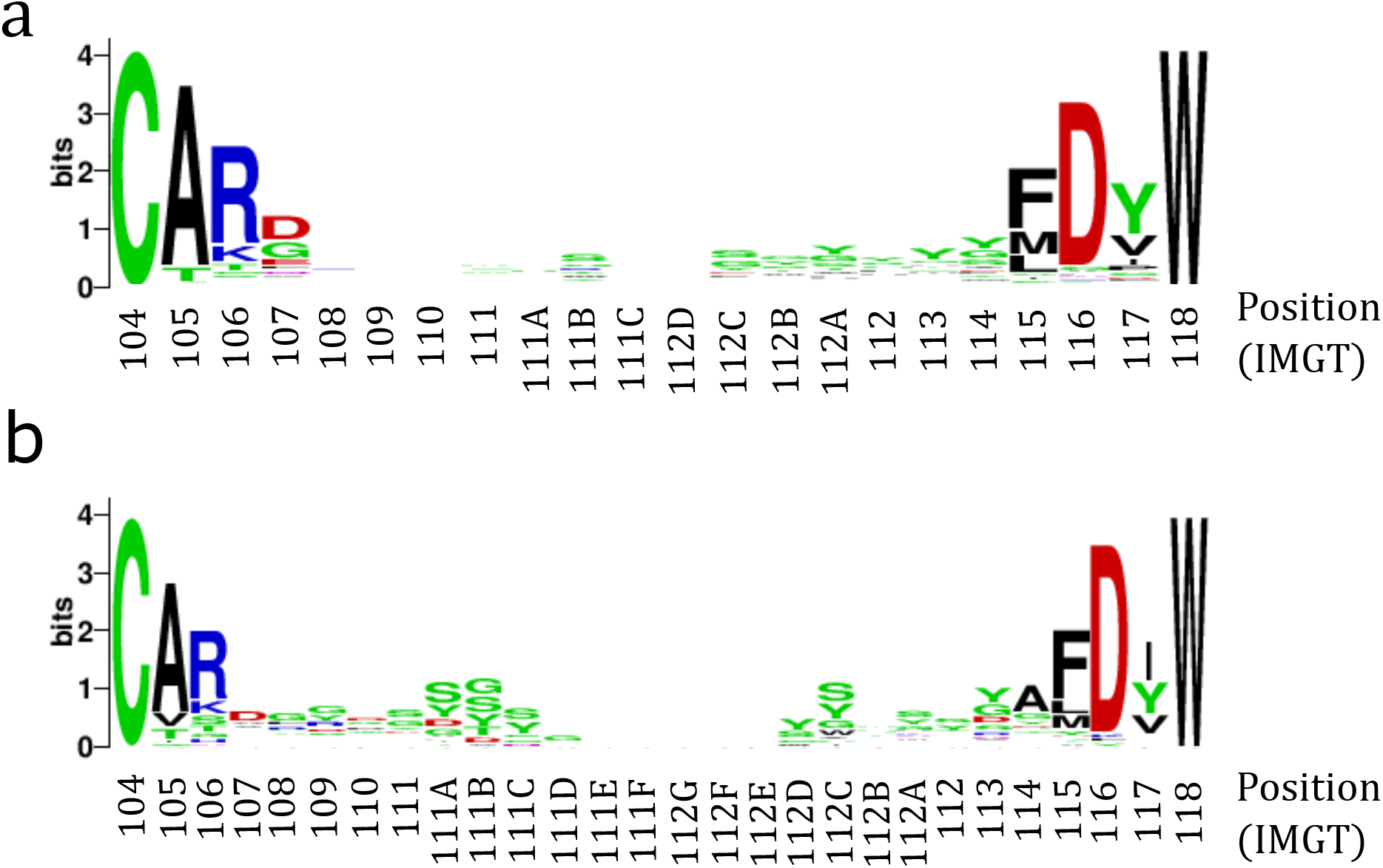
WebLogo (73) plots showing the entropy and distribution of residues at each IMGT (20) CDRH3 position for (a) SARS-CoV Receptor Binding Domain (RBD) binding antibodies, and (b) MERS-CoV RBD binding antibodies.

Finally, we evaluated the closest sequence identity match between all SARS-CoV binding CDRH3s and the over 500 million CDRH3s in our Observed Antibody Space (OAS) database (78). The OAS database is a regularly updated project to catalogue all publicly available immune repertoire sequencing experiments (currently over 60 studies), providing cleaned amino acid sequence datasets binned by individual and other useful metadata. We assume that the vast majority of this sampled population is serologically naive to SARS-CoV-1 and SARS-CoV-2, given both the high infection rate and that there is currently no evidence to suggest that exposure to common cold coronaviruses yields SARS-CoV cross-reactive antibodies (79). It follows that the presence of CDRH3s shown to bind SARS-CoV but that have high sequence identity matches to OAS may be less useful for diagnosing SARS-CoV-2 exposure. Table 1 contains all CDRH3s for which we obtained 100% identity matches to a CDRH3 in OAS. A table showing all maximum sequence identity matches is available as Supplementary File 1 and the full dataset (with OAS metadata for matches) is available as Supplementary File 2.

**Table 1.**
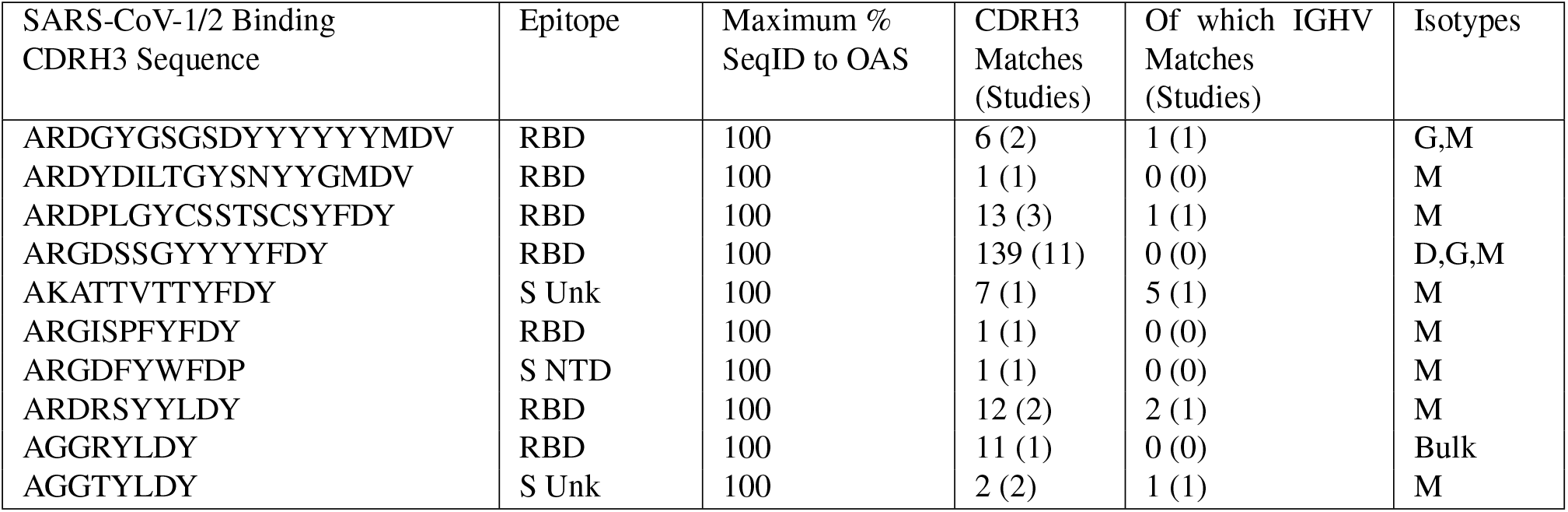
The ten SARS-CoV binding antibody CDRH3s from CoV-AbDab that matched with 100% sequence identity to a CDRH3 sequence in the OAS database. A full table showing all CDRH3s with their closest matches to an OAS sequence is available as SI Table 1. RBD = Receptor-Binding Domain, Spike protein, SeqID = Sequence Identity, OAS = Observed Antibody Space database (78), Unk = Unknown.

We observed that 10/69 (14.4%) SARS-CoV binding CDRH3s had 100% sequence matches to at least one sequence in OAS, while 45/69 (65.2%) had at least one 80% or greater sequence identity match. The mean sequence identity match was 83%. Interestingly, two of the CDRH3s with 100% matches (ARDPLGYCSSTSCSYFDY: 3C7/5E10/6B1, ARGDSSGYYYYFDY: S304) were found to be proximal to sequences isolated in the recent Stanford SARS-CoV-2 patient serum investigation (26). Exact clonal matches (V gene + high CDRH3 identity) were considerably rarer, implying full clonotyping may need to be performed on SARS-CoV-2 repertoires in order to identify genuine responding antibodies. Conversely, some CDRH3s from SARS-CoV-2 neutralising antibodies found in SARS-CoV-1 (mAb S309 (6)) and SARS-CoV-2 (mAb 32D4 (25)) responding repertoires have considerably lower than average closest sequence identity matches to OAS (70% and 67% respectively).

## Community Contributions

We have attempted to identify all existing published information on SARS-CoV and MERS-CoV binding antibodies, however encourage users to inform us of any historical investigations we may have missed. We are also reaching out to authors of new studies characterising coronavirus binding antibodies to send us their data in Excel or CSV format. Data and queries may be sent to us by email. Minimum requirements for addition to our database are the full antibody/nanobody variable domain sequence, binding or neutralising data for at least one specified coronavirus protein, and a link to a relevant preprint, publication, or patent. Through these submissions and our own efforts to track the scientific literature, we hope to provide a central community resource for coronavirus antibody sequence and structural information.

## Usage

Currently, the database can be queried by a search term (e.g. SARS-CoV-2) and ordered by any metadata field for maximum interpretability. Users can download the entire database as a CSV file and bulk download all ANARCI numberings, IMGT-numbered PDB files, and IMGT-numbered homology models.

## Accessibility

CoV-AbDab is free to access and download without registration and is hosted at *http://opig.stats.ox.ac.uk/webapps/coronavirus*.

## Patents

CoV-AbDab uses the following patents as a primary source of antibody/nanobody sequences: CN1664100, CN1903878, CN100374464, CN104447986, CN106380517, EP2112164, KR101828794, KR101969696, KR20190122283, KR20200020411, US7396914, WO2005/012360, WO2005/054469, WO2005/060520, WO2006/095180, WO2008/035894, WO2015/179535, WO2016/138160, and WO2019039891.

## ACKNOWLEDGEMENTS

This work was supported by an Engineering and Physical Sciences Research Council (EPSRC) and Medical Research Council (MRC) grant [EP/L016044/1] awarded to MIJR, and Biotechnology and Biological Sciences Research Council (BBSRC) grant [BB/M011224/1] award to AK, and funding from GlaxoSmithKline plc, UCB Pharma Ltd., AstraZeneca plc, and F. Hoffmann-La Roche. We would like to thank the many authors who have responded in a timely and helpful manner to our requests for data.

## Notes

http://opig.stats.ox.ac.uk/webapps/coronavirus

### Competing Interest Statement

The authors have declared no competing interest.

