## Supplementary Information for "CoV-AbDab: the Coronavirus Antibody Database"

### Supporting Information
#### CoV-AbDab: the Coronavirus Antibody Database

Raybould MIJ, Kovaltsuk A, Marks C, and Deane CM

Corresponding Author: Charlotte M. Deane


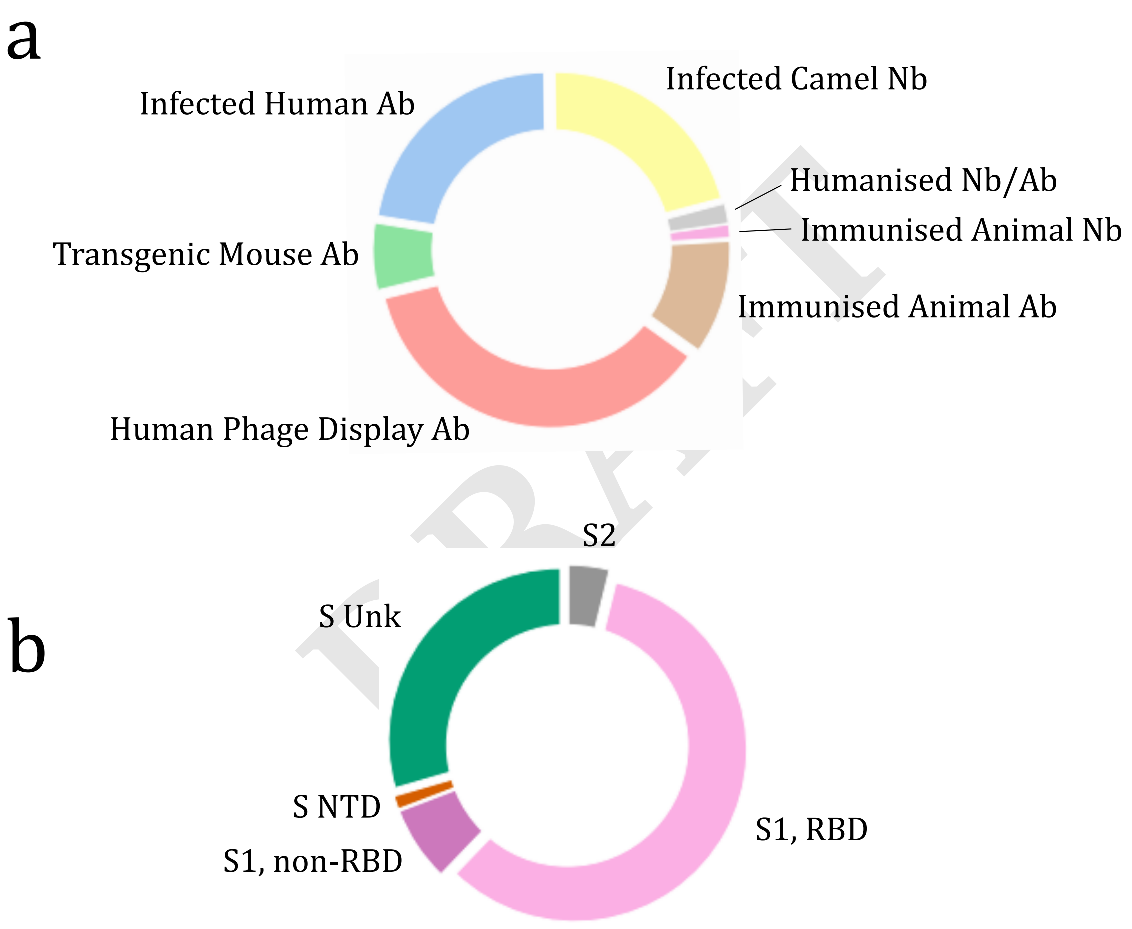


**Fig. 1.** Donut charts showing (a) the origins of all identified MERS-CoV binders and (b) the protein targets of identified MERS-CoV binders. Spike protein binders are further classified by targeted domain. S = Spike protein, Unk = Unknown, NTD = N-Terminal Domain, RBD = Receptor Binding Domain, S1 = Spike protein S1 domain, S2 = Spike protein S2 domain.

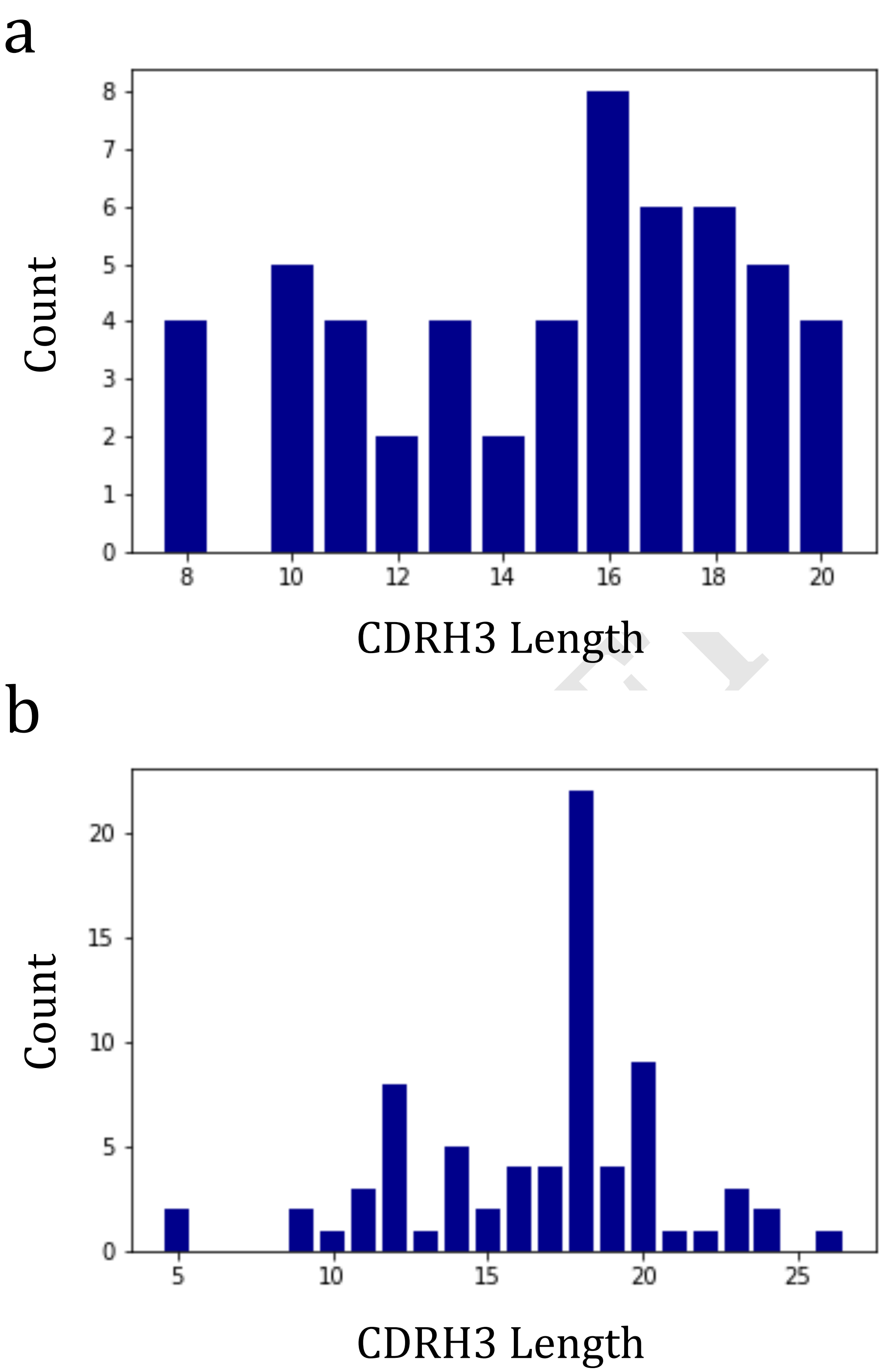


**Fig. 2.** Graphs of CDRH3 length frequencies for (a) SARS-CoV-1/2 Receptor Binding Domain (RBD) binding antibodies and (b) MERS-CoV RBD binding antibodies.

#### Files

The following files are hosted at *http://opig.stats.ox.ac.uk/resources*:

1. *Table_SARS-CoV_CDRH3s_inOAS.csv*

A table summarising the closest OAS sequence identity matches to each SARS-CoV binding CDRH3 in CoV-AbDab.

2. *Dataset_SARS-CoV_CDRH3s_inOAS.pkl*

The full dataset of closest OAS sequence identity matches to each SARS-CoV binding CDRH3 in CoV-AbDab. Contains full metadata for each OAS match (e.g. full VH sequence, study, individual, disease state, *etc*).

DRAFT
